# Rat hepatitis E virus (HEV) cross-species infection and transmission in pigs

**DOI:** 10.1101/2023.07.06.547957

**Authors:** Kush K. Yadav, Patricia A Boley, Carolyn M Lee, Saroj Khatiwada, Kwonil Jung, Thamonpan Laocharoensuk, Jake Hofstetter, Ronna Wood, Juliette Hanson, Scott P. Kenney

**Affiliations:** Center for Food Animal Health, Department of Animal Sciences, The Ohio State University, Wooster, Ohio, USA; Department of Veterinary Preventive Medicine, The Ohio State University, College of Veterinary Medicine, Columbus, Ohio, USA

**Keywords:** Rat, hepatitis E, virus, pigs, infectious, zoonotic, model

## Abstract

Rocahepevirus ratti, an emerging hepatitis E virus (HEV), has recently been found to be infectious to humans. Rats are a primary reservoir of the virus; thus, it is referred to as “rat HEV”. Rats are often found on swine farms in close contact with pigs. Our goal was to determine whether swine may serve as a transmission host for rat HEV by characterizing an infectious cDNA clone of a zoonotic rat HEV, strain LCK-3110, in vitro and in vivo. RNA transcripts of LCK-3110 were constructed and assessed for their replicative capacity in cell culture and in gnotobiotic pigs. Fecal suspension from rat HEV-positive gnotobiotic pigs was inoculated into conventional pigs cohoused with naïve pigs. Our results demonstrated that capped RNA transcripts of LCK-3110 rat HEV replicated in vitro and successfully infected conventional pigs that transmit the virus to cohoused animals. The infectious clone of rat HEV may afford an opportunity to study the genetic mechanisms of rat HEV cross-species infection and tissue tropism.

**Significance Statement:** New zoonotic strains of Rocahepevirus ratti (rat HEV) have emerged infecting both immunocompetent and immunosuppressed people through unknown transmission sources. Pigs are a primary source of transmission for human HEV strains and could be serving a similar role for rat HEV transmission as rats are a common pest found on swine farms worldwide. Rats could be transmitting rat HEV to pigs which could then be transmitted to humans. Determining susceptibility of pigs to emerging zoonotic rat HEV strains can define potentially new transmission routes to inform public health policy and could provide pathology models for rat HEV disease.

## Introduction

Hepatitis E is a disease caused by hepatitis E virus (HEV) (1). HEV is a widespread pathogen found in wild animals (boar, deer) (2, 3), domestic pigs (4, 5), rats (6, 7), chickens (8, 9), and humans (10, 11). The disease in humans is mostly defined as self-limiting but can be lethal in people with underlying health conditions (12, 13), immunosuppressed (14, 15), and during pregnancy (16-19). HEV infection is increasingly being found as a complication for solid organ transplant patients (15, 20-22) where HEV can persist in immunosuppressed individuals, leading to liver cirrhosis (23).

HEV is classified in the family Hepeviridae that comprises two subfamilies: Orthohepevirinae and Parahepevirinae (24). Out of 4 genera (Paslahepevirus, Avihepevirus, Rocahepevirus and Chirohepevirus) in the Orthohepevirinae, Paslahepevirus is known to cause clinical disease in humans with all 8 genotypes potentially capable of infecting humans (gt1 through gt8) (Fig. 1) (19). Gt1 and gt2 are restricted to humans, whereas HEV gt3 through gt8 are zoonotically transmitted from pigs and other species to humans (25). The most common spillover strains are gt3 and gt4 which act as foodborne illnesses transmitting from primarily pork products to humans (10, 26-29). Multiple studies have demonstrated the presence of HEV in pig slaughter houses (30-32), pork products at grocery stores (33-37). Zoonotic transmission of pig gt3 HEV has been a concern for immunosuppressed patients as they develop chronic disease. The clinical outcomes appear to be becoming more severe with more frequent extrahepatic disease manifestations (20, 38, 39).

**Figure 1.**
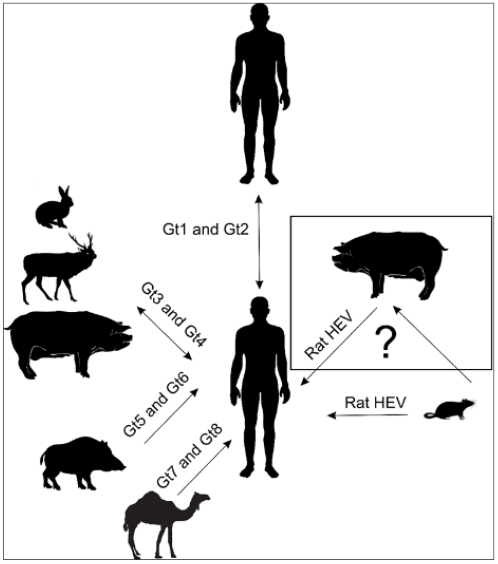
HEV transmission pathways to humans. Paslahepevirus (gt1 and gt2) has been demonstrated to obligately transmit between humans. Paslahepevirus (gt3 to gt8) has been considered zoonotic to humans. Rat HEV has been seen to cause infection in humans via the unknown source of transmission.

Rocahepevirus ratti (rat HEV), formerly Orthohepevirus C, circulates at a high level in rat populations (40-46). Initial experiments in 2013 with rat HEV concluded it to be non-infectious to humans as evidenced by the absence of viral replication in non-human primates (NHPs) (47). Infections in humans were not noted until 2018 when it was first reported in an immunosuppressed human case in Hong Kong (48). Since this initial case, rat HEV has been shown to be infectious to both immunosuppressed (48-51) and immunocompetent humans (52). Rat HEV disease has been defined by mild hepatitis but has been demonstrated to progress to persistent infection in immunosuppressed patients (49). Rat HEV is an international concern as it has been detected in a traveler from Africa to Canada, suggesting a global prevalence of this emerging understudied virus (52). Experimental cross-species transmission studies of rat HEV (V-105) to rhesus and cynomolgus monkeys demonstrated fecal shedding and seroconversion with no significant alteration in liver enzymes (53). Rat HEV recovered from cynomolgus and rhesus monkeys successfully infected both nude and Sprague-Dawley rats (53). Such findings of the cross-species transmission of rat HEV necessitate understanding of the rat HEV host range, transmission to agriculturally important species, and pathogenesis in detail.

There are several areas of concern with emerging rat HEV cases in humans. Serological presence of IgG antibodies against Paslahepevirus did not provide protection against rat HEV (48, 51). Commonly used RT-qPCR screening assays are based on Paslahepevirus, thus it has been unable to differentially detect rat HEV (48). Retrospective studies to find the transmission source of rat HEV could not identify the exact transmission route of rat HEV to humans (48). Pigs are known as one of the main reservoirs (54) and transmission sources of HEV to humans (55). This emphasizes the need to understand the relevance of this agriculturally important species as a potential intermediate source of rat HEV transmission to humans. Genetic analyses revealed that all human cases of rat HEV infection reported in Hong Kong are genetically close to LCK-3110 strain (51). In this study, we successfully developed a rat HEV infectious clone (LCK-3110) and demonstrated its ability to develop infection in pigs.

## Results

### 3.1 Construction of Full-Length cDNA Clone of Rat HEV LCK-3110 Strain and Determination of its Replicative Ability in Cell Lines from Multiple Species

A full-length genomic cDNA clone of rat HEV LCK-3110 strain was constructed and cloned into the pSP64 poly (A) vector (Fig. 2A). The full-length cDNA clone (pSP64-rat HEV-poly (A)) was sequenced, restriction enzyme digested, and run through agarose gel electrophoresis to check the rat HEV insertion, 6,941 base pairs (Fig. 2B).

**Figure 2.**
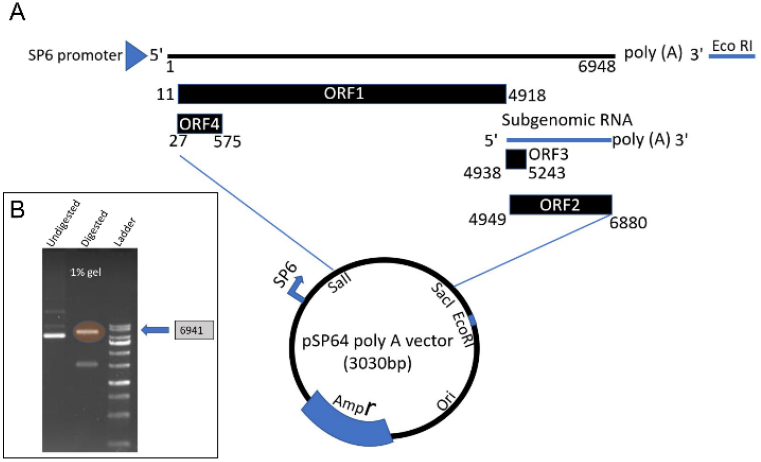
Successful insertion of rat HEV into pSP64 poly A vector. (A) Schematic representation of full-length genomic rat HEV insertion in the pSP64 poly A vector. A cDNA clone of rat HEV LCK-3110 strain was constructed and cloned into the pSP64 poly (A) vector using unique restriction sites, SaII and SacI. It contains EcoRI restriction sites at the 3’ end of the viral genome. (B) Results from 1% gel electrophoresis demonstrating the insertion of the full-length rat HEV.

To test the infectivity of the pSP64-rat HEV-poly (A) cDNA clone in vitro, capped viral RNAs were transcribed in vitro (Fig. 3A). The capped RNAs were transfected into human hepatoma (Huh7), mouse subcutaneous tissue (LMTK), human carcinoma lung tissue (A549) and swine testicular (ST) cells to determine its replication ability. Cells were fixed and probed for ORF2 expression via an immunofluorescence assay (IFA). As ORF2 is translated from subgenomic mRNA, it is synthesized only during the later stages of HEV replication, serving as an indicator of complete viral replication (56, 57). To demonstrate whether HEV transcripts have productively replicated in the target cells, we assessed events at the single-cell level using IFA and flow cytometry. Employing antibodies (rabbit anti-ORF2 polyclonal serum) directed against Paslahepevirus ORF2 capsid protein, we detected cells expressing the ORF2 protein suggesting replication (Fig. 3C). Total numbers of ORF2 expressing cells were quantified by IFA against ORF2 coupled with flow cytometry. As depicted in Fig. 3B, ST, A549, and LMTK cells were more permissive for rat HEV replication than Huh7 cells. Approximately, 8%, 6.5%, 7%, and 4% ORF2-positive cells were observed in ST, LMTK, A549, and Huh7 cells, respectively. Additionally, Huh7 cells were transfected with the cell culture adapted Kernow C1 P6 gt3 HEV strain and two non-cell culture adapted strains (Kernow C1 P1; gt3 and Sar55; gt1) to build a cutoff point to demonstrate the ORF2 percent positive cells. Cell supernatant and cell lysates harvested on day 5 demonstrated viral RNA loads in both cell lysate and supernatants from LMTK, A549, ST, and Huh7 cells via RT-qPCR (Fig. 3D, 3E). These results demonstrate that the pSP64-rat HEV-poly (A) cDNA clone can produce RNA which is replication competent in vitro.

**Figure 3.**
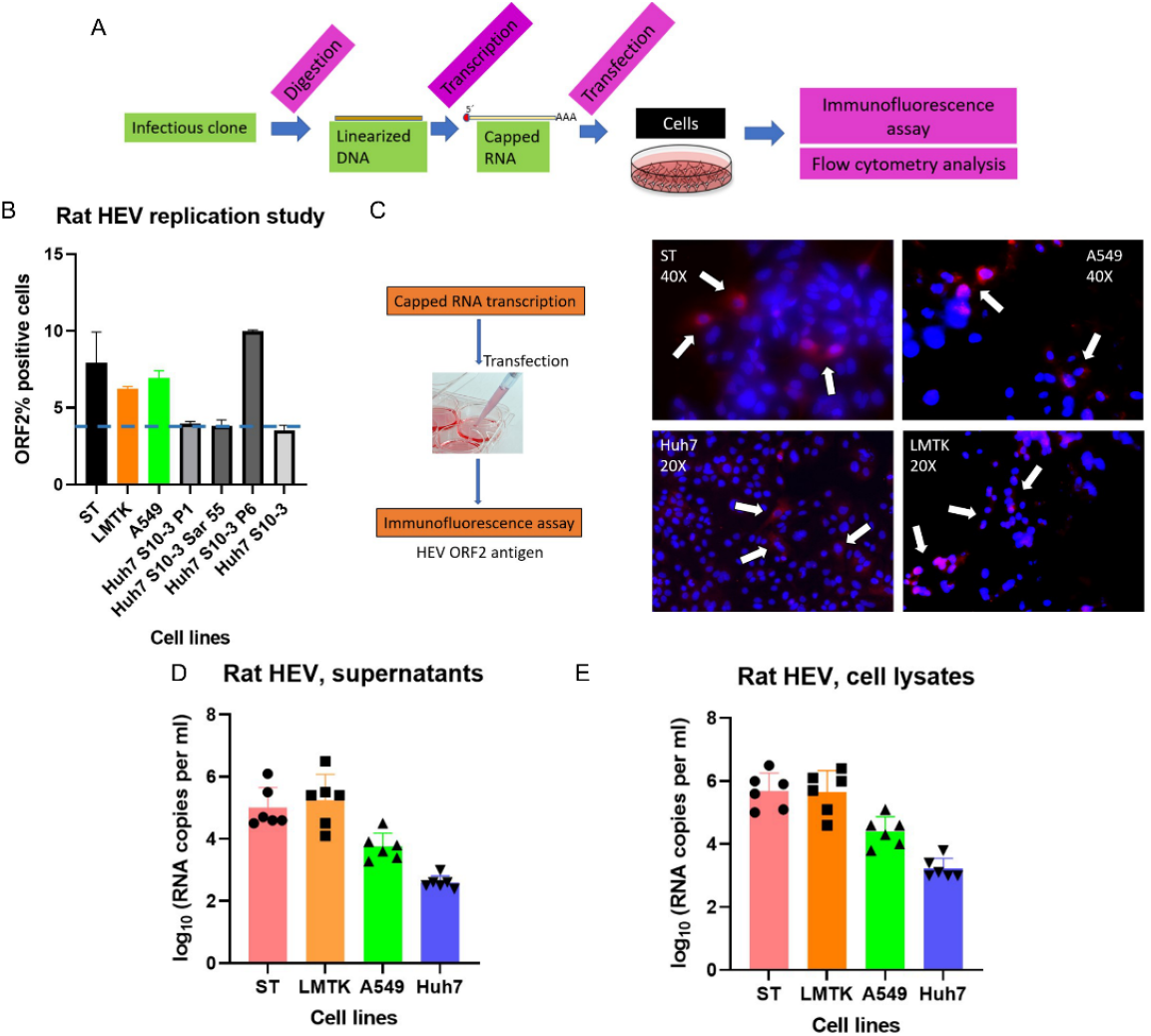
Rat HEV is replication competent in cells. (A) Workflow of HEV capping and transfection of target cells. LMTK, A549, Huh7, ST cell lines were transfected with in vitro transcribed capped HEV RNA (rat HEV). (B) Flow cytometry quantification of LMTK, A549, Huh7, ST cells transfected with capped RNA transcripts of rat HEV; Huh7 transfected with cell culture adaptive Kernow-C1 genotype 3 P6 strain and non-cell culture adaptive genotype 1 Sar55 and Kernow-C1 genotype 3 P1 strain. Sar55, P6 and P1 belong to Paslahepevirus is used as the control to determine the replication ability of rat HEV. The assay was performed in the cells harvested on day 5 post transfection. Samples were fixed in methanol and probed with rabbit anti-ORF2 followed by goat anti-rabbit alexa fluor 594 antibodies. Each bar (mean ± SD) represents separate transfections stained in parallel and displays the mean of two independent biological experiments with three replicates per sample. (C) Immunofluorescence detection of HEV ORF2 antigen in methanol fixed LMTK, A549, Huh7, ST cells after 5 days post transfection. Cells are stained with goat anti-rabbit IgG H&L combined with anti-rabbit Alexa fluor 594 (red), and 4’
s, 6-diamidino-2-phenylindole (DAPI) (blue). (D) RT-qPCR data from the supernatants collected from day 5 of replication assay. (E) RT-qPCR data from the cell lysates collected from day 5 of replication assay.

### 3.2 Capped RNA Transcripts of the Rat HEV LCK-3110 were Infectious when Intrahepatically Injected into the Livers of Gnotobiotic Pigs

Since high titer HEV infectious virus is difficult to obtain in vitro, infectivity or pathogenicity studies with live infectious virus are limited and sometimes results in less interpretable data due to the low starting titer of available infectious virus (58). With the successful construction of an infectious cDNA clone of rat HEV LCK-3110 strain, we bypassed cell culture virus propagation and amplified the virus directly in animals via an intrahepatic inoculation procedure (59) with capped RNA transcripts. The above-described intrahepatic inoculation procedure (also referred to as in vivo transfection) has been successfully used for pathogenicity studies of swine and human HEV (59-62). Gnotobiotic pigs were intrahepatically inoculated with the LCK3110 capped viral RNA and tested for evidence of rat HEV infection in the pigs (Fig. 4A). Rat HEV was detected in fecal material starting at day 9 post inoculation (Fig. 4C). Viral RNA in feces and serum was detected until the study termination date, 35 days post inoculation (dpi). Rat HEV RNA was also detected in bile, liver, spleen, ileum, kidney, urinary bladder, urine, feces, brain, and cerebrospinal fluid (CSF) (Fig. 4D) at 35 dpi. As a positive control, gnotobiotic pigs were inoculated intravenously (IV) with a 10% fecal suspension from US-2 HEV infected pigs (kindly provided by Dr. XJ Meng, Virginia Tech) (Fig. 4B). Evidence of US-2 HEV infection, including fecal virus shedding, was also detected (Fig. 4C).

**Figure 4.**
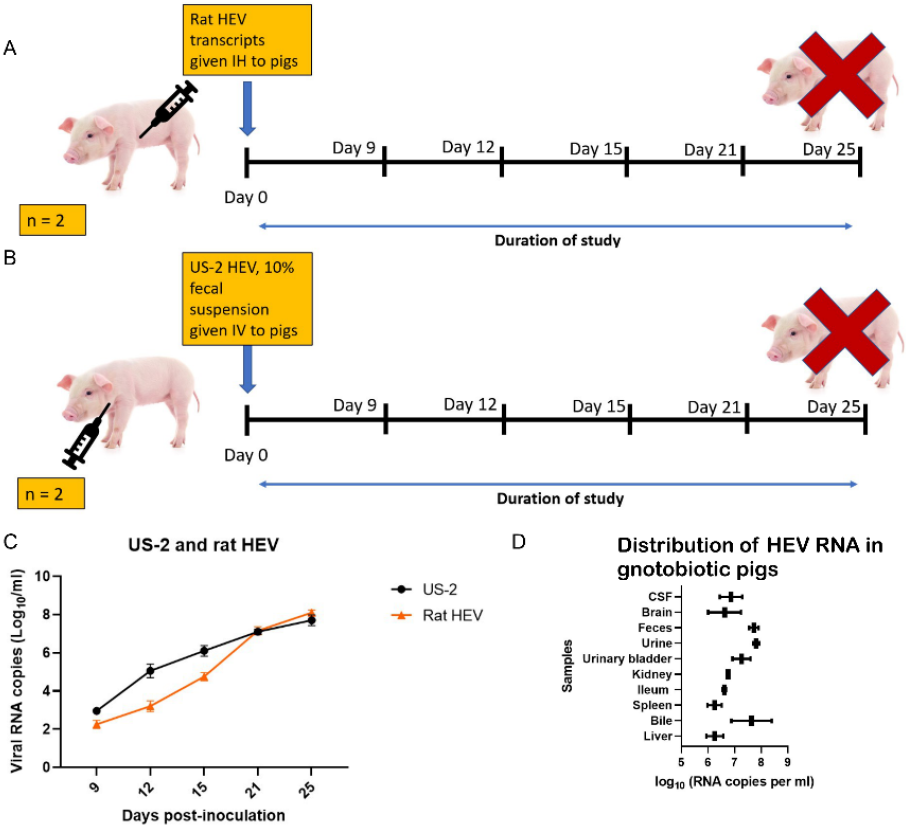
Successful amplification of rat HEV and US-2 HEV in pigs. (A) Schematic representation of the experimental design. Capped rat HEV transcripts inoculated intrahepatically (IH) to pigs. (B) Schematic representation of the experimental design. Fecal suspension (10%) harvested from US-2 positive pigs inoculated intravenously (IV) to pigs. (C) Fecal viral shedding of rat HEV and US-2 HEV. (D) HEV RNA loads in CSF, brain, feces, urine, urinary bladder, kidney, ileum, spleen, bile, liver in rat HEV inoculated pigs.

### 3.3 Characterization of the Pathogenicity of Rat HEV LCK-3110 Recovered from Gnotobiotic Pigs in Conventional Pigs

Intestinal contents derived from the intrahepatically inoculated gnotobiotic pigs on day 35 was used for the preparation of an infectious stock used in the pathogenicity study. Group A (n=3) was injected IV with rat HEV (2 × 108), group B (n=2) was injected IV with US-2 HEV (2 × 10^8^) and group C (n=2) was IV injected with PBS. Rat HEV RNA and US-2 HEV RNA were detected in feces (Fig. 5A) and sera (Fig. 5B) by RT-qPCR in group A and group B with primers specific for rat HEV strain and US-2 HEV strain, respectively. Fecal virus shedding and viremia were detected from 1 week post inoculation in both experimental groups (A and B) with variable rates of detection and rat HEV inoculated pigs shedding more than the US-2 HEV (Fig. 5). Rat HEV produced a higher fecal titer in pigs at week 1 post infection than US-2. US-2 fecal shedding appeared to taper off beginning at 3 weeks post infection, whereas the rat HEV infected pigs continued to increase fecal shedding through the end of the study, 35 dpi. Sentinel pigs in the rat HEV infected room began to shed virus in their feces at week 2 post comingling and appeared to begin tapering off at week 4 post comingling (Fig. 5A).

**Figure 5.**
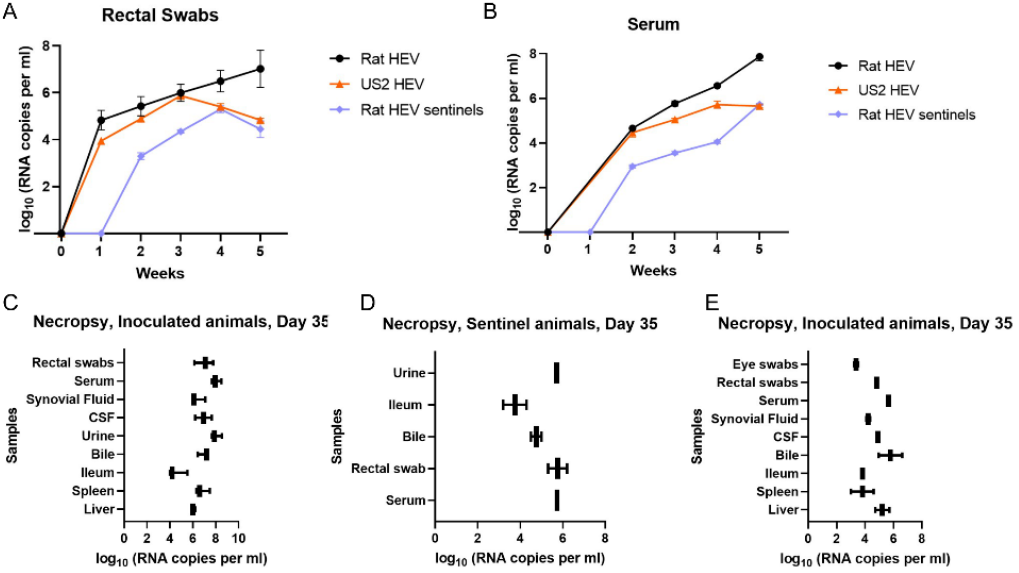
Assessment of the conventional pig model for genotype 1 rat HEV via intravenous inoculation of 10% fecal suspension harvested from gnotobiotic pigs. (A) Viral RNA in feces was determined by RT-qPCR between groups. (B) Viral RNA in serum was determined by RT-qPCR between groups. (C) HEV RNA loads in rectal swabs, serum, synovial fluid, CSF, urine, bile, ileum, spleen, liver in rat HEV inoculated pigs. (D) HEV RNA loads in urine, Ileum, Bile, Rectal swab, serum in sentinel pigs comingled with rat HEV infected pigs. (E) HEV RNA loads in rectal swabs, serum, synovial fluid, CSF, urine, bile, ileum, spleen, liver in US-2 HEV inoculated pigs.

Trends in viremia mimicked fecal shedding with rat HEV RNA in the serum increasing for the duration of the study, US-2 RNA peaked at around week 4 post infection, and rat HEV sentinel animal RNA titers increased for the duration of the study (Fig. 5B). Total antibodies against HEV were examined in all groups from day 0, 14, 21, 28, and 35. Interestingly, pigs inoculated with rat HEV did not seroconvert (Fig. 7F) but US-2 HEV inoculated pigs seroconverted by day 21 (Fig. 7E). Rat HEV sentinels did not seroconvert (Fig. 7G) similar to rat HEV inoculated pigs. Negative control pigs did not seroconvert throughout the study (Fig. 7D).

In the rat HEV infected group, all pigs necropsied on 35 dpi had viral RNA in feces and serum demonstrating fecal viral shedding and viremia. Other samples, from necropsy such as liver, spleen, and ileum were also positive for viral RNA. Bodily fluids such as bile, urine, CSF, and synovial fluid were positive for rat HEV RNA with bile, urine, and feces having the highest concentration of RNA (Fig. 5C).

In the US-2 HEV infected group, all pigs necropsied at 35 dpi had viral RNA in feces and serum (Fig. 5A, B). Other samples, such as liver, spleen, ileum, bile, CSF, synovial fluid, eye swabs were positive for US-2 HEV (Fig. 5E). Mock challenged animals remained negative in fecal shedding and viremia.

In HEV-inoculated pigs, histopathological lesions were limited to the liver. Both rat HEV and US-2 HEV-inoculated pigs had moderate, multifocal lymphohistiocytic hepatitis at 35 dpi (Fig 6A). Compared with rat HEV-inoculated pigs, the sentinel pigs had less infiltration of non-suppurative inflammatory cells, demonstrating milder hepatitis or early status of inflammatory responses to HEV infection (Fig. 6B). By IHC, HEV ORF2 protein was identified in the livers of rat HEV and US-2 HEV-inoculated pigs. The HEV ORF2 protein was seen mainly in the cytoplasm or perinuclear region of hepatocytes at 35 dpi (Fig. 6C). Mock-infected pigs occasionally had mild splenic or hepatic congestion, but they did not exhibit HEV ORF2 protein in the tissues tested.

**Figure 6.**
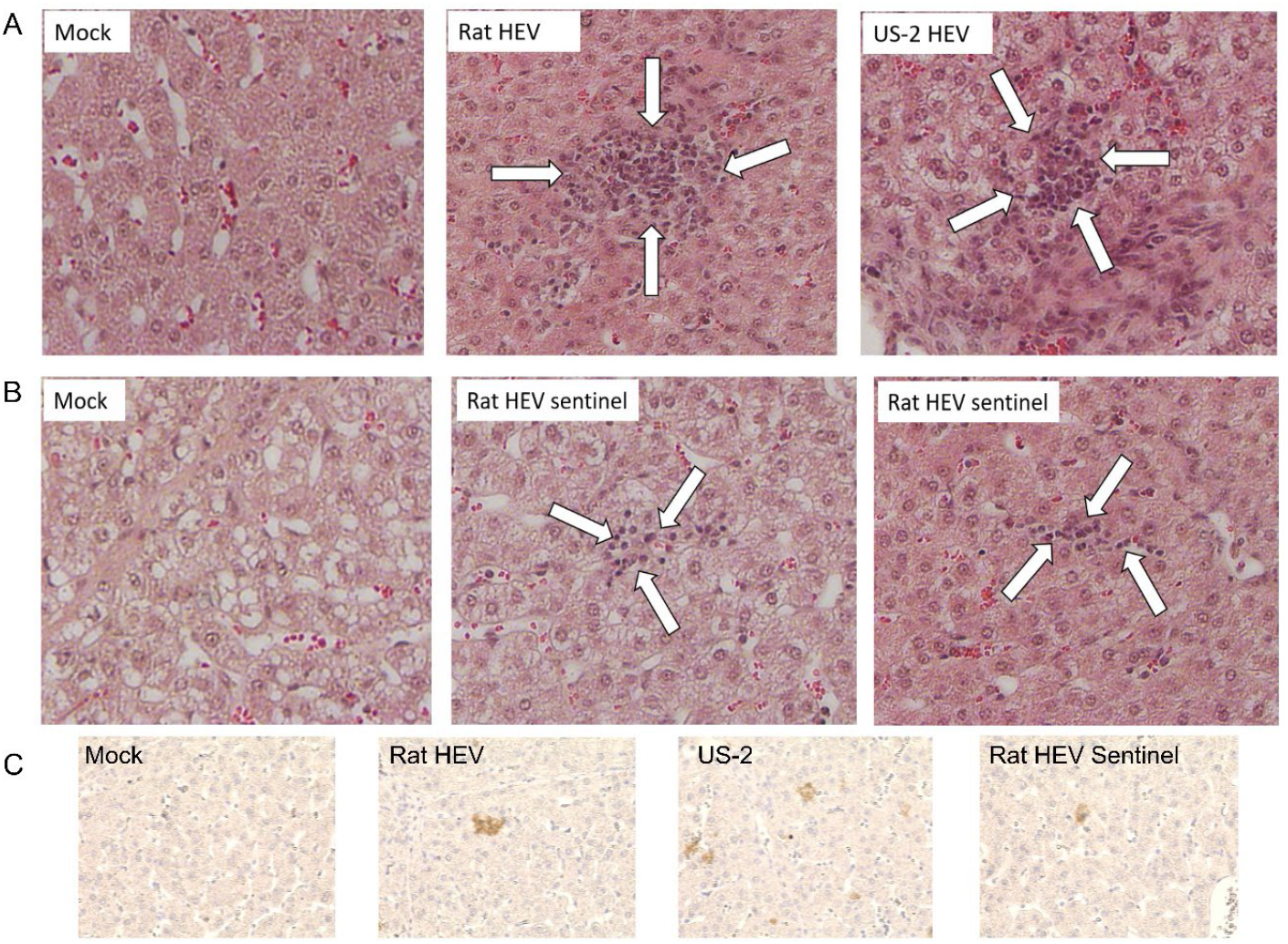
Histopathology and immunohistochemistry (IHC) in the liver of pigs inoculated with rat HEV or US-2 HEV and the sentinel pigs at 35 days post-infection (dpi). (A) Hematoxylin and eosin (H&E)-stained liver of a rat HEV-inoculated pig, showing moderate lymphohistiocytic hepatitis (arrows). Note that US-2 HEV-inoculated and mock pigs exhibit moderate lymphohistiocytic hepatitis (arrows) and no lesions, respectively. (B) H&E-stained liver of sentinel rat HEV group pigs, showing mild lymphocytic or lymphohistiocytic hepatitis (arrows). Note that the mock pig shows no lesions. (C) IHC-stained liver of a rat HEV-inoculated pig, showing a small amount of HEV ORF2 protein (brown stain). Note that US-2 HEV-inoculated and mock pigs exhibit a small amount of HEV ORF2 protein (brown stain) and no IHC-positive cells, respectively. The rat HEV sentinel pig also has a small amount of HEV ORF2 protein in the liver. Chromogenic detection of HEV ORF2 protein via 3,3’
s-Diaminobenzidine (DAB) staining. Original magnification (A-C), all x400. H&E or IHC-stained images were taken by Keyence BZ-X800.

The liver enzyme tests indicated no significant difference between the groups in ALT and AST during the period of infection, suggesting that no serious liver damage was induced in the pigs by rat HEV infection (Fig. 7A, 7B). The levels of GGT were higher in rat HEV inoculated pigs by day 35 indicating the initiation of cholestasis in pigs (Fig. 7C).

**Figure 7.**
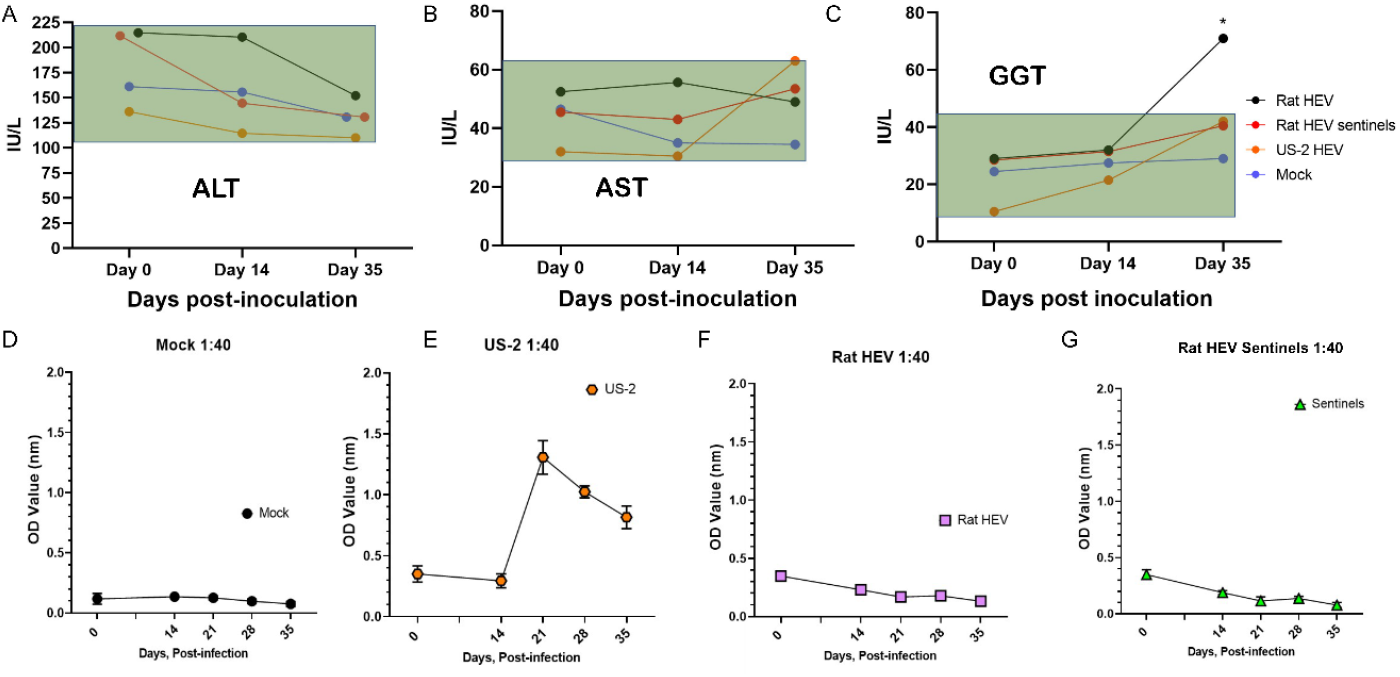
Liver function enzyme levels and seroconversion to HEV during rat HEV infection. The serum biochemistry of alanine aminotransferase (ALT) (A), aspartate aminotransferase (AST) (B), and gamma-glutamyl transferase (GGT) (C) was examined in rat HEV infected pigs. The values inside the green block represent the normal expected range of the enzymes in pigs. Increase in the GGT by day 35 is seen in the pigs inoculated with the rat HEV. All values are shown in units per liter (IU/L). (D) Negative control pigs did not show any seroconversion by day 35. (E) Positive control pigs (US-2 HEV inoculated), the titer of the anti-HEV antibody increased after day 14. (F, G) No seroconversion was seen in rat HEV inoculated pigs and rat HEV sentinel pigs.

### 3.4 Rat HEV Transmission Through Fecal-Oral Route to Sentinel Pigs

The fecal-oral route is considered as one of the major routes of transmission of HEV. To study whether the virus is efficiently transmitted to sentinel pigs, naïve animals were co-housed with pigs that had been infected with the fecal inoculum. Sentinel pigs added to the rat HEV inoculated group on 7 dpi started shedding virus in feces after 7 days post contact (Fig. 5A). Viremia (Fig. 5B) was evident in the sentinel pigs as early as 7 days post contact (Fig. 5B). High titers of viral RNA were detected in the feces and blood of these animals, although these titers were lower than those of the originally infected pigs. However, only bile, urine, and ileum demonstrated viral RNA after 28 days post contact with the rat HEV inoculated pigs (Fig. 5D). Overall, these data confirm the efficient transmission of rat HEV between co-housed animals.

### 3.5 Human Hepatoma Cell Lines Huh7 and HepG2/C3A Support LCK-3110 Replication

Huh7 and HepG2/C3A were inoculated with LCK-3110 rat HEV obtained from intestinal contents of rat HEV inoculated conventional pigs. We detected rat HEV RNA in supernatants from both cell lines (Fig. 8A) inoculated with pig intestinal contents on day 2, 4, and 6. RNA detected on day 0 (6 hours post inoculation) was considered as background with attachment of the virus to the cell surfaces. RNA loads in cell lysates collected on day 6 demonstrated successful viral cell entry and replication (Fig. 8A). IFA (Fig. 8B) of Huh7 and HepG2/C3A cells on day 6 post inoculation confirmed the presence of ORF2 protein in the inoculated cells. Replication in Huh7 S10-3 cells was comparatively better than HepG2/C3A cells (Fig. 8A). These results demonstrate the ability of rat HEV derived from conventional pig feces to replicate in human hepatoma cell lines.

**Figure 8.**
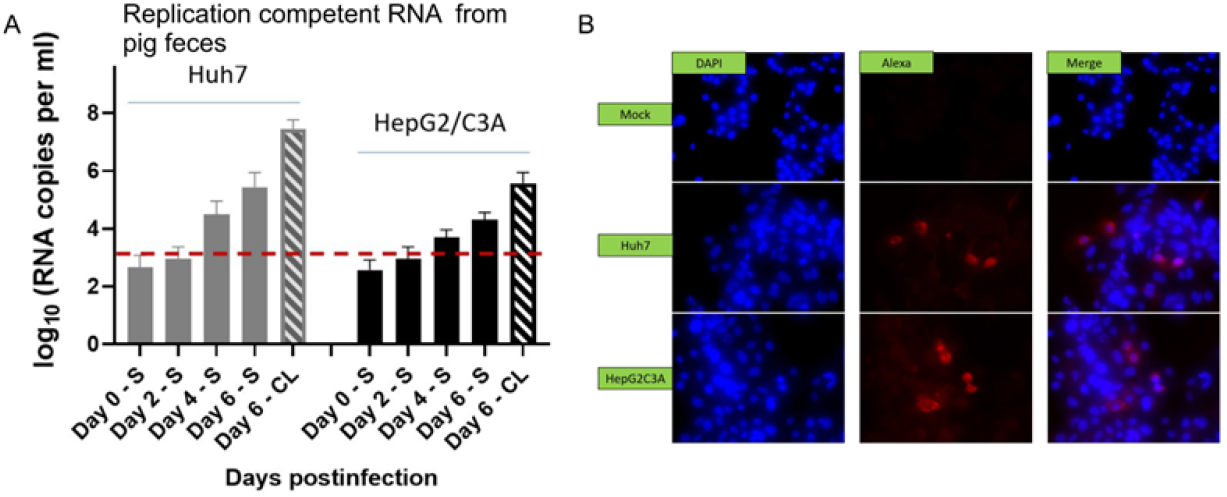
Isolation of rat HEV from infected pig feces in cell culture. (A) Rat HEV RNA loads in culture supernatant (S) and cell lysates (CL) of Huh7 and HepG2/C3A cell lines after inoculation by filtered fecal suspension from rat HEV inoculated pigs. Independent biological experiments, mean ± SD of 4 replicates, are presented. Red line represents the cut-off value demonstrating the background referring to the attachments of the virus to the cell surfaces (B) Immunofluorescence detection of HEV ORF2 antigen in methanol fixed Huh7 and HepG2/C3A cells after 6 days post inoculation. Cells are stained with goat anti-rabbit IgG H&L combined with anti-rabbit Alexa fluor 594 (red), and 4’
s, 6-diamidino-2-phenylindole (DAPI) (blue).

## Discussion

Increasing HEV host range is a topic of considerable concern (63). Historically, all HEV infections in humans have been attributed to Paslahepevirus (gt1 to gt8), but recently Rocahepevirus ratti has been identified as a causative agent in human disease (48-51). In 2013, rat HEV strain LA-B350 isolated from rats in US (GenBank: KM516906.1) was considered non-infectious to humans based on the genetic diversity from Paslahepevirus strains and its inability to infect non-human primates and pigs (47, 64). However, from 2017 to 2022, thirteen hepatitis E cases attributed to rat HEV have been identified, and to date most of them were in Hong Kong. Genetic analyses indicated that these strains were close to LCK-3110 strain (GenBank: MG813927.1) (51). In our study, we developed the LCK-3110 infectious clone and explored its infectious ability to known primary reservoirs of Paslahepevirus HEV, namely pigs. Our results reveal that pigs can serve as an intermediate host for rat HEV infection in humans and should be studied further for this transmission potential.

Other recent studies determining rat HEV cross species transmission potential showed the V-105 strain of Rocahepevirus ratti isolated from Vietnam was capable of infecting primates (53). Strain V-105 is genetically closest to the LCK-3110 strain group and shares 93.7% nucleotide identity with LCK-3110. In our study, we demonstrate that the LCK-3110 strain can infect pigs producing suppurative hepatitis and shedding virus with no significant alterations in liver enzymes as seen in cynomolgus and rhesus monkeys with V-105 (53).

Rat HEV has been detected in rats from Japan (6), United States (47), Germany (65, 66), Indonesia (46), Vietnam (67) and many other countries, suggesting a geographically widespread distribution of rat HEV among rat populations. Although non-persistent infection has been described in rats with rat HEV (68), higher rates of progression to persistent rat HEV infection have been reported in immunosuppressed humans even after reductions in immunosuppressive drug therapy (50). This finding is concerning and needs to be further studied to understand how to best treat rat HEV infection in immunosuppressed humans utilizing in vivo models. Multiple studies describing rat HEV in humans have reported no contact of the patients with rats (48, 49). Retrospective studies to identify the transmission source of rat HEV to humans failed to demonstrate the presence of rat HEV in the organs of local rats, from swab samples of drains or in rat fecal samples. Only one rat’s internal organs harvested in 2012 were reported positive for rat HEV (48). This emphasizes that there could be an intermediate source for rat HEV infection that is mediating the transmission of the virus to humans.

Some studies have investigated commensal rats and pigs in proximity to the residence of rat HEV positive humans (49), which might have contributed to infection. Only 7/159 R. norvegicus reported positive for rat HEV (49). All 172 pig rectal swabs were negative for rat HEV RNA (49). The findings do not definitively eliminate pigs as a transmission source due to the low level and transient nature of HEV RNA shed and inability to serologically separate rat HEV from Paslahepevirus HEV strains. Pigs are well known reservoirs of Paslahepevirus HEV strains without producing major clinical signs and could complicate serosurveillance studies. Time of sample collection and viral load in the commensal rats are the limitations during RNA surveillance studies. Another study demonstrated two R. rattus that were positive for both rat HEV and genotype 3 swine HEV. Swine HEV detected in rats was identical to the swine HEV in pigs. The detection of HEV in the liver is an indication of active viral replication, as the liver is the major organ of viral replication (69). But swine HEV was not detected in the liver of rats. The presence could be attributed to the ingestion of feces from swine pens but raises the question of possible transmission of HEV strains between the two species. Rats and pigs can easily be exposed to each other’s feces containing high amounts of virus. Thus, the probability of pigs being infected with rat HEV via contaminated feces cannot be eliminated. With our experimental finding that pigs can be infected with rat HEV, it further enhances our knowledge of rat HEV host range, necessitating further study on this potential transmission vector. More surveillance and the development of rat HEV specific serological tools are needed to appropriately address pigs as potential transmission vectors for rat HEV.

Our study demonstrates the presence of viral RNA in the cerebrospinal fluid (CSF) of rat HEV infected pigs. The recent discovery of HEV invasion of microvascular endothelial cells and the ability to cross the blood-brain barrier invading the nervous system demands the need to study rat HEV effects in the central nervous system (70). Meningoencephalitis followed by death have been reported in an immunosuppressed organ transplant recipient with persistent rat HEV infection. Examination of the CSF revealed the presence of rat HEV (49). Investigations of rat HEV antigenicity showed that rat HEV is highly divergent from Paslahepevirus that reduces the ability of defined diagnostic assays against HEV. Failure of cross protective antibodies against rat HEV even when previously exposed to Paslahepevirus necessitates an update to the existing diagnostic assays for HEV (51). Amino acid 455-603 of ORF2 refers to the protruding domain which is the immunodominant epitope in HEV. The average intergenotypic amino acid identity within Paslahepevirus is 89.5% while the aa identity with LCK-3110 was only 48% (51). Lack of monoclonal antibody (mAb) cross-binding renders HEV antigen kits ineffective for rat HEV diagnosis. The reduced sensitivity and specificity of the Paslahepevirus ORF2 could be the reason for the lack of seroconversion seen in our rat HEV inoculated pigs. However, difference in seroconversion has been demonstrated in pigs with various HEV strains. Pigs experimentally infected with swine HEV gt3 seroconverted in 55 days whereas pigs infected with US-2 HEV seroconverted within a week (71). We found similar results with US-2 HEV but there was no seroconversion seen in the rat HEV group despite viral shedding in feces, viremia, and presence of HEV in the liver. We speculate that the time allocated for the study was possibly not sufficient to demonstrate seroconversion in the rat HEV inoculated pigs. RT-qPCR assays designed for rat HEV detecting rat HEV in an immunosuppressed patient who tested negative for Paslahepevirus HEV is a great example demonstrating the failure of current conventional hepatitis E molecular diagnostics (50). We did not see any major elevation of liver enzymes such as Alanine aminotransferase (ALT) and aspartate aminotransferase (AST) in the US-2 infected pigs validating previous findings (71). ALT and AST levels in the rat HEV inoculated pigs were no different than US-2 HEV and mock pigs. However, gamma-glutamyl transferase (GGT) levels were significantly higher in rat HEV inoculated group. Increased GGT activity is an indicator of cholestasis in pigs (72) and is used in the investigation of hepato-pancreatic or renal disorders (73). Rat HEV positive asymptomatic human patients were described with very mild liver dysfunction matching our results in the animal model (50).

In conclusion, we have constructed an infectious cDNA clone of a genotype 1 LCK-3110 strain of rat HEV and demonstrated its infectivity in vitro and in vivo. We produced infectious viruses through intrahepatic RNA inoculation of gnotobiotic pigs and characterized the pathogenicity of rat HEV through IV injection of intestinal contents to conventional pigs. Like Paslahepevirus HEV infection in pigs, gross lesions were not detected with only microscopic lesions evident in the liver of rat HEV infected pigs. Furthermore, we revealed that pigs could be an intermediate source of emerging zoonotic rat HEV transmission to humans.

## Materials and Methods

### Construction of full-length cDNA clone of rat HEV

The full-length genomic sequence of the LCK-3110 strain of rat HEV (Genbank MG813927.1) was artificially synthesized (Genscript). The plasmid pSP64 poly (A) vector (Promega) was used to clone the full-length rat HEV genome between the SalI and SacI sites (Fig 2A). After the successful insertion, the plasmid was transformed into stable E. Coli (NEB) and grown overnight at 37°C in the presence of ampicillin.

### Linearization of Plasmid DNA

For linearization, plasmid DNA encoding rat HEV was linearized using EcoRI (NEB). Five percent of the reaction was subjected to gel electrophoresis with ethidium bromide staining and visualized with ultraviolet light to verify that linearization had occurred.

### In Vitro Transcription for IFA

Viral capped mRNA (rat HEV) was made from linearized DNA using the Promega Ribomax Large Scale RNA Production System SP6 (Promega PRP 1300) and ARCA CAP (TriLink Biotechnologies). The fidelity of transcripts was assessed and normalized by agarose gel electrophoresis.

### Cell Culture

LMTK (isolated from mouse subcutaneous tissue, ATCC:CCL-1.3), A549 (isolated from human lung carcinoma, ATCC:CRM-CCL-185), Huh7 (isolated from human liver carcinoma), ST (isolated from swine testicular, ATCC:CRL-1746) and Huh7 (human hepatoma cells) S10-3 subclone (74, 75) were used for the studies. LMTK, Huh7 S10-3, and ST cells were cultured in DMEM (Dulbecco Modified Eagle’s Medium) containing 10% FBS (Fetal bovine serum). A549 cells were cultured in F12/K media containing 10% FBS.

### Transfection of ST, LMTK, Huh7, and A549 cells

RNA transfection was achieved using Mirus Trans-IT mRNA transfection kit. After 48 hours of transfection, cells were passaged 1:3 to three new wells and cells were incubated for an additional 3 days.

### Flow cytometry of In Vitro-Transcribed Capped Rocahepevirus ratti HEV RNA Transfected (ST, Huh7, LMTK and A549) Cells

Five days post transfection (dpt), cells were trypsinized and pelleted. Cells were then fixed in 200 μL of 100% methanol at 4°C. After overnight storage at -80°C, cells were centrifuged out of methanol, washed, and resuspended in phosphate buffered saline (PBS). Cells were blocked-in blocking solution (5% non-fat dried milk, 0.1% Triton X-100 in PBS; PBST) in a 96-well plate for 30 min at 37°C. Cells were then washed with PBS once before probing with primary antibody— rabbit anti truncated ORF2 HEV (57, 64) diluted 1:50 in blocking solution for 30 min at 37°C. After washing twice in PBS, cells were incubated with secondary antibody-goat anti-rabbit-phycoerythrin (PE) (Life Technologies) diluted to 1:200 PBS for 30 min at 37°C. Cells were then washed twice in PBS, resuspended in 200 µL of PBS. Fluorescence was analyzed for 100,000 events using a flow cytometer (BD Accuri C6 Plus, Biosciences, San Diego, CA, USA). Gates were set to exclude dead cells, doublet discrimination based on forward and side scatter profiles, and mock infected cells were used to gate background fluorescence (75).

### Indirect Immunofluorescence

At 5 dpt, transfected cells were fixed in 100% cold methanol, permeabilized with PBST, and blocked with 5% non-fat milk (Sigma-Aldrich, St. Louis, MO, USA). Immunostaining of ORF2-encoded capsid protein was performed using a 1:200 blocking buffer diluted rabbit anti truncated ORF2 HEV antibody for 30 minutes at 37°C. Cells were washed 3 times with PBST. A fluorescently labeled goat anti-rabbit IgG H&L antibody (Alexa Fluor 594; abcam, Cambridge, FL, USA) was used at a dilution of 1:400 in PBS to detect bound primary antibodies. 4′,6-diamidino-2-phenylindole (DAPI) was used to stain the nucleus. For quantification of virus infectivity, wells were manually observed with a fluorescent microscope (Keyence) for specific fluorescence, and the presence of fluorescent foci was recorded. A fluorescent focus was defined as a minimum of one to two cells showing clear intracytoplasmic fluorescence (75).

### Intrahepatic Inoculation of Capped RNA Transcripts from Rat HEV Infectious cDNA Clone

All animal experiments in this study were approved by The Ohio State University Institutional Animal Care and Use Committee. Four gnotobiotic pigs from 1 sow were derived near term and maintained in two different sterile isolation units (two/unit) as described previously (76). On day 2, pigs were administered capped rat HEV transcripts via intrahepatic injection. The other two pigs were intravenously inoculated with 10% fecal suspension from US-2 positive pigs (positive control). Rectal swabs from pigs were collected on days 9, 12, 15, 21, and 25. Pigs were humanely euthanized on day 25. Blood, intestinal contents, tissues (brain, urinary bladder, kidney, duodenum, jejunum, ileum, pancreas, liver, bile, spleen) and bodily fluids (cerebrospinal fluid, urine, bile) were harvested from rat HEV inoculated pigs.

### Intravenous Inoculation of 10% Fecal Suspension of Rat HEV in Conventional Pigs

Ten conventional pigs, negative for rat HEV and US-2 HEV RNA in rectal swabs were randomly divided into three groups (A, B, C). Group A (n=3) was intravenously administered 10% fecal suspension derived from rat HEV inoculated gnotobiotic pigs. Group B (n=2) was intravenously administered 10% fecal suspension derived from US-2 HEV inoculated gnotobiotic pigs. Group C (n=2) was intravenously administered PBS. After a week, two pigs negative for rat HEV were added as sentinels with Group A (rat HEV). Rectal swabbing was done weekly in all groups. Blood collection was done on day 0, 14, 21, 28, and 35. Pigs were humanely euthanized on day 35. Blood, intestinal contents, eye swabs, tissues (brain, urinary bladder, kidney, duodenum, jejunum, ileum, pancreas, liver, bile, spleen, lungs, skeletal muscle) and bodily fluids (cerebrospinal fluid, urine, bile) were harvested from the pigs.

### RNA Extraction and RT-qPCR

RNA extraction was performed using trizol reagent (Invitrogen) from harvested cell supernatant and cell lysates on day 5 from A549, LMTK, Huh7 and ST cells. Similarly, RNA was extracted from serum, blood, eye swabs, homogenized tissues, and bodily fluids. Reverse transcriptase quantitative polymerase chain reaction (RT-qPCR) was performed. A one step RT-qPCR was carried out using TaqMan Fast Virus 1-step Master Mix (ThermoScietific) under a protocol of 50°C for 15 min, 95°C for 2 min and 45 cycles of 95 for 5 seconds and 60°C for 30 seconds (Mastercycler RealPlex). A forward primer rat HEV F, 5’-CTTGTTGAGCTYTTCTCCCCT-3’, a reverse primer, 5’-CTGTACCGGATGCGACCAA-3’, and a probe 5’-FAM-TGCAGCTTGTCTTTGARCCC -Dabcyl-3’ were used for the detection of rat HEV. A 10-fold serial dilution of the capped rat HEV RNA (10^7^ to 10^1^ copies) was used as the standard for the quantification of the viral genome copy numbers. A similar procedure was done for pig samples experimentally infected with US-2 HEV. A forward primer US-2 HEV F, 5′-GGTGGTTTCTGGGGTGAC-3′, a reverse primer, 5′-AGGGGTTGGTTGGATGAA-3′, and a probe 5’-FAM-TGATTCTCAGCCCTTCGC-Dabcyl-3’ were used for the detection of US-2 HEV. A 10-fold serial dilution of the capped US-2 HEV RNA (10^7^ to 10^1^ copies) was used as the standard for the quantification of the viral genome copy numbers.

### Histological Examinations and Immunohistochemistry (IHC)

The liver (from each lobe), pancreas, duodenum, jejunum, ileum, and spleen of virus- or mock-infected pigs were collected and fixed in 10% neutral buffered formalin. Tissues were embedded, sectioned (3.5 µm), and stained with Gill’s hematoxylin and eosin (H&E) for light microscopic examination as described previously (77). Formalin-fixed tissue sections were tested by IHC for the detection of HEV, as previously described with slight modifications (78). The rabbit anti-HEV-ORF2 antibody was used as the primary antibody and a horseradish peroxidase conjugated anti-rabbit antibody (BioGeneX) was used for visualization as brown staining. Stained tissues were counterstained with hematoxylin.

### Recombinant Proteins

Two nucleotide sequences encoding for amino acids 391-620 of gt3 ORF2 were codon optimized for bacterial expression and synthesized commercially (IDT), inserted into bacterial T7 expression vector pRSETa (Invitrogen), and expressed. Recombinant protein was produced using BL21 (DE3) chemically competent cells via autoinduction. Proteins were analyzed via SDS-PAGE and western blot. The bacteria were lysed with B-Per™ reagent (Thermofisher) (5 mL/gram) with 1 mM ethylenediaminetetraacetic acid (EDTA). ELISA protein was solubilized and purified with Ni-NTA columns followed by dialysis. Protein was quantified using Bradford assay.

### ELISA

ELISAs were modified and optimized to detect HEV ORF2-specific IgG antibodies in serum. Five µg/ml of purified plasmid lysate diluted in carbonate buffer (20 mM Na2CO3, 20 mM NaHCO3, pH 9.6) were bound to Nunc Maxisorp 96 well plates (Thermofisher) at 50 µl per well at 4°C overnight. The following morning, after washing, 150 µl of blocking buffer [4% nonfat dried milk in PBS with 0.1% Tween (PBST 0.1%)] was added to the antigen-coated wells and incubated for 2 hours at 37°C. Plates were washed with PBS and 50 µl of serum was heat inactivated at 56°C for 30 minutes. Inactivated serum was 2 -fold serially diluted in blocking buffer and added to each well. The plates were incubated for 1 hour at 37°C. After washing, 50 µl of HRP-conjugated secondary antibody [anti-swine (Sigma)] in 4% NFDM/PBST (0.1%) at a dilution of 1:200K was added and incubated at 37°C for 1 h. Wells were washed with PBST (0.1%) five times between each step. 3,3’,5,5’-tetramethylbenzidine (TMB) substrate (Seracare) was added and incubated for approximately 10 minutes, and the reaction was stopped by adding 50 µl of 0.3 mol/L sulfuric acid. Plates were read at an absorbance of 450 nm using a SpectraMax F5 plate reader (Molecular Devices). All experiments were done under the same conditions with each sample tested three times.

### Liver Enzyme Levels

ALT, AST, and GGT values in the pig’s sera were monitored using VETSCAN VS2 (Zoetis). The geometric mean titer of ALT, AST, and GGT in negative control animals was considered the normal ALT titer, and a fold greater or more was considered as alteration in the liver enzymes.

### Virus Culture

Huh7 S10-3 and HepG2/C3A cells were seeded in six well plates at 2 × 10^5^ cells per well in 2 ml of DMEM with 10%FBS and penicillin (100 units/ml) and streptomycin (100g/ml) and incubated at 37°C for 24 hours. The infections were performed utilizing a 10% fecal suspension diluted 1:5 in DMEM and 0.45μm filtered. Culture media was removed, and cells were inoculated with 1 ml of the resulting solution (1 × 10^7^ viral RNA copies). At room temperature, plates were rocked for 1 hour and then incubated at 37°C for 6 hours. The inoculum was removed, and fresh culture media was added. Supernatants were collected on day 0, 2, 4, and 6. Supernatants and lysates were tested for rat HEV via RT-qPCR.

### Statistical Analyses and Reproducibility

All quantitative data are presented with the mean and standard deviation. Analyses of independent data were performed by Student’s unpaired two-tailed t test. Statistical analyses were carried out using GraphPad Prism 9.4.1. p < 0.05 was considered significant.

## Acknowledgments

We thank Megan Strother and Sara Tallmadge at Center for Food Animal Health, Department of Animal Sciences, The Ohio State University, Wooster, Ohio, USA for their assistance in animal care.

## References

1. Khuroo MS (2011) Discovery of hepatitis E: the epidemic non-A, non-B hepatitis 30 years down the memory lane. Virus Res 161(1):3–14.

2. Ferri G, Piccinini A, Olivastri A, & Vergara A (2022) Hepatitis E virus detection in hunted wild boar (Sus scrofa) livers in Central Italy. Italian journal of food safety 11(2):9979.

3. Boadella M (2015) Hepatitis E in wild ungulates: A review. Small Ruminant Research 128:64–71.

4. Amorim AR, Mendes GS, Pena GPA, & Santos N (2018) Hepatitis E virus infection of slaughtered healthy pigs in Brazil. Zoonoses and public health 65(5):501–504.

5. Lee HS, et al. (2020) Prevalence and phylogenetic analysis of hepatitis E virus in pigs in Vietnam. BMC Vet Res 16(1):333.

6. Takahashi M, et al. (2022) First detection and characterization of rat hepatitis E Virus (HEV-C1) in Japan. Virus Res 314:198766.

7. De Sabato L, et al. (2020) Detection of hepatitis E virus RNA in rats caught in pig farms from Northern Italy. Zoonoses and public health 67(1):62–69.

8. Meng K, Liu M, Zhang Y, Yuan X, & Xu H (2022) A genetically novel avian Hepatitis E virus in China. Virus Genes 58(6):589–593.

9. Siedlecka M, Kublicka A, Wieliczko A, & Matczuk AK (2022) Molecular detection of avian hepatitis E virus (Orthohepevirus B) in chickens, ducks, geese, and western capercaillies in Poland. PLoS One 17(6):e0269854.

10. Treagus S, Wright C, Baker-Austin C, Longdon B, & Lowther J (2021) The Foodborne Transmission of Hepatitis E Virus to Humans. Food and environmental virology 13(2):127–145.

11. Whitsett M, Feldman DM, & Jacobson I (2020) Hepatitis E Virus Infection in the United States: Current Understanding of the Prevalence and Significance in the Liver Transplant Patient Population and Proposed Diagnostic and Treatment Strategies. Liver transplantation : official publication of the American Association for the Study of Liver Diseases and the International Liver Transplantation Society 26(5):709–717.

12. Zhang L, et al. (2017) Prevalence of hepatitis E virus infection among blood donors in mainland China: a meta-analysis. Transfusion 57(2):248–257.

13. Wang L, et al. (2020) Absence of hepatitis E virus RNA in semen samples of infertile male in China. Gut 69(7):1363.

14. Yadav KK & Kenney SP (2021) Hepatitis E Virus Immunopathogenesis. Pathogens 10(9).

15. Damiris K, Aghaie Meybodi M, Niazi M, & Pyrsopoulos N (2022) Hepatitis E in immunocompromised individuals. World journal of hepatology 14(3):482–494.

16. Tissera G, et al. (2020) Hepatitis E virus infection in pregnant women, Argentina. BMC Infect Dis 20(1):368.

17. Ismail MB, et al. (2020) Seroprevalence of hepatitis E virus in pregnant women in northern Lebanon. Eastern Mediterranean health journal = La revue de sante de la Mediterranee orientale = al-Majallah al-sihhiyah li-sharq al-mutawassit 26(5):580–585.

18. Zhang F, et al. (2022) Clinical features of sporadic hepatitis E virus infection in pregnant women in Shanghai, China. The Journal of infection 84(1):64–70.

19. Yadav KK & Kenney SP (2022) Hepatitis E Virus Zoonotic Axis. Zoonoses: Infections Affecting Humans and Animals, ed Sing A (Springer International Publishing, Cham), pp 1–28.

20. Kamar N, et al. (2008) Hepatitis E virus and chronic hepatitis in organ-transplant recipients. The New England journal of medicine 358(8):811–817.

21. Fang SY & Han H (2017) Hepatitis E viral infection in solid organ transplant patients. Curr Opin Organ Transplant 22(4):351–355.

22. de Niet A, et al. (2012) Chronic hepatitis E after solid organ transplantation. Neth J Med 70(6):261–266.

23. Kamar N & Pischke S (2019) Acute and Persistent Hepatitis E Virus Genotype 3 and 4 Infection: Clinical Features, Pathogenesis, and Treatment. Cold Spring Harbor perspectives in medicine 9(7).

24. Purdy MA, et al. (2022) ICTV virus taxonomy profile: Hepeviridae 2022. Journal of General Virology 103(9):001778.

25. Sooryanarain H & Meng XJ (2020) Swine hepatitis E virus: Cross-species infection, pork safety and chronic infection. Virus research 284:197985.

26. Cossaboom CM, et al. (2016) Risk factors and sources of foodborne hepatitis E virus infection in the United States. J Med Virol 88(9):1641–1645.

27. Dai X, et al. (2013) Hepatitis E virus genotype 4, Nanjing, China, 2001-2011. Emerg Infect Dis 19(9):1528–1530.

28. Cossaboom CM, Córdoba L, Dryman BA, & Meng XJ (2011) Hepatitis E virus in rabbits, Virginia, USA. Emerging infectious diseases 17(11):2047–2049.

29. Feagins AR, et al. (2011) Intergenotypic chimeric hepatitis E viruses (HEVs) with the genotype 4 human HEV capsid gene in the backbone of genotype 3 swine HEV are infectious in pigs. Virus research 156(1-2):141–146.

30. Chelli E, et al. (2021) Hepatitis E Virus Occurrence in Pigs Slaughtered in Italy. Animals (Basel) 11(2).

31. Boxman ILA, et al. (2022) High prevalence of acute hepatitis E virus infection in pigs in Dutch slaughterhouses. International journal of food microbiology 379:109830.

32. Sooryanarain H, et al. (2020) Hepatitis E Virus in Pigs from Slaughterhouses, United States, 2017-2019. Emerging infectious diseases 26(2):354–357.

33. Pallerla SR, et al. (2021) Hepatitis E virus genome detection in commercial pork livers and pork meat products in Germany. Journal of viral hepatitis 28(1):196–204.

34. Gutiérrez-Vergara C, Quintero J, Duarte JF, Suescún JP, & López-Herrera A (2015) Detection of hepatitis E virus genome in pig livers in Antioquia, Colombia. Genet Mol Res 14(1):2890–2899.

35. Soares VM, et al. (2022) Detection of adenovirus, rotavirus, and hepatitis E virus in meat cuts marketed in Uruguaiana, Rio Grande do Sul, Brazil. One health (Amsterdam, Netherlands) 14:100377.

36. Harrison L, Ramos TM, Wu X, & DiCaprio E (2021) Presence of hepatitis E virus in commercially available pork products. International journal of food microbiology 339:109033.

37. Ruggeri FM, et al. (2013) Zoonotic transmission of hepatitis E virus in industrialized countries. The new microbiologica 36(4):331–344.

38. Alric L, Bonnet D, Laurent G, Kamar N, & Izopet J (2010) Chronic Hepatitis E Virus Infection: Successful Virologic Response to Pegylated Interferon-α Therapy. Annals of Internal Medicine 153(2):135–136.

39. Dalton HR, Bendall RP, Keane FE, Tedder RS, & Ijaz S (2009) Persistent carriage of hepatitis E virus in patients with HIV infection. N Engl J Med 361(10):1025–1027.

40. Kanai Y, et al. (2012) Hepatitis E virus in Norway rats (Rattus norvegicus) captured around a pig farm. BMC research notes 5:4.

41. Ryll R, et al. (2017) Detection of rat hepatitis E virus in wild Norway rats (Rattus norvegicus) and Black rats (Rattus rattus) from 11 European countries. Vet Microbiol 208:58–68.

42. Johne R, et al. (2012) Rat hepatitis E virus: geographical clustering within Germany and serological detection in wild Norway rats (Rattus norvegicus). Infection, genetics and evolution : journal of molecular epidemiology and evolutionary genetics in infectious diseases 12(5):947–956.

43. Simanavicius M, et al. (2018) Detection of rat hepatitis E virus, but not human pathogenic hepatitis E virus genotype 1-4 infections in wild rats from Lithuania. Vet Microbiol 221:129–133.

44. Murphy EG, et al. (2019) First detection of Hepatitis E virus (Orthohepevirus C) in wild brown rats (Rattus norvegicus) from Great Britain. Zoonoses and public health 66(6):686–694.

45. Lack JB, Volk K, & Van Den Bussche RA (2012) Hepatitis E virus genotype 3 in wild rats, United States. Emerg Infect Dis 18(8):1268–1273.

46. Mulyanto, et al. (2013) Frequent detection and characterization of hepatitis E virus variants in wild rats (Rattus rattus) in Indonesia. Archives of virology 158(1):87–96.

47. Purcell RH, et al. (2011) Hepatitis E virus in rats, Los Angeles, California, USA. Emerg Infect Dis 17(12):2216–2222.

48. Sridhar S, et al. (2018) Rat Hepatitis E Virus as Cause of Persistent Hepatitis after Liver Transplant. Emerg Infect Dis 24(12):2241–2250.

49. Sridhar S, et al. (2021) Transmission of Rat Hepatitis E Virus Infection to Humans in Hong Kong: A Clinical and Epidemiological Analysis. Hepatology (Baltimore, Md.) 73(1):10–22.

50. Sridhar S, et al. (2022) Hepatitis E Virus Species C Infection in Humans, Hong Kong. Clin Infect Dis 75(2):288–296.

51. Sridhar S, et al. (2021) Multimodal investigation of rat hepatitis E virus antigenicity: Implications for infection, diagnostics, and vaccine efficacy. Journal of hepatology 74(6):1315–1324.

52. Andonov A, et al. (2019) Rat Hepatitis E Virus Linked to Severe Acute Hepatitis in an Immunocompetent Patient. J Infect Dis 220(6):951–955.

53. Yang F, et al. (2022) Experimental Cross-Species Transmission of Rat Hepatitis E Virus to Rhesus and Cynomolgus Monkeys. Viruses 14(2).

54. Meng XJ (2011) From barnyard to food table: the omnipresence of hepatitis E virus and risk for zoonotic infection and food safety. Virus Res 161(1):23–30.

55. Johne R, Althof N, Nöckler K, & Falkenhagen A (2022) [Hepatitis E virus-a zoonotic virus: distribution, transmission pathways, and relevance for food safety]. Bundesgesundheitsblatt, Gesundheitsforschung, Gesundheitsschutz 65(2):202–208.

56. Graff J, Torian U, Nguyen H, & Emerson SU (2006) A bicistronic subgenomic mRNA encodes both the ORF2 and ORF3 proteins of hepatitis E virus. Journal of virology 80(12):5919–5926.

57. Kenney SP & Meng XJ (2015) The lysine residues within the human ribosomal protein S17 sequence naturally inserted into the viral nonstructural protein of a unique strain of hepatitis E virus are important for enhanced virus replication. Journal of virology 89(7):3793–3803.

58. Kwon HM, et al. (2011) Construction of an infectious cDNA clone of avian hepatitis E virus (avian HEV) recovered from a clinically healthy chicken in the United States and characterization of its pathogenicity in specific-pathogen-free chickens. Veterinary microbiology 147(3-4)310–319.

59. Huang YW, et al. (2005) Capped RNA Transcripts of Full-Length cDNA Clones of Swine Hepatitis E Virus Are Replication Competent When Transfected into Huh7 Cells and Infectious When Intrahepatically Inoculated into Pigs. Journal of virology 79(3):1552.

60. Emerson SU, et al. (2001) Recombinant hepatitis E virus genomes infectious for primates: importance of capping and discovery of a cis-reactive element. Proc Natl Acad Sci U S A 98(26):15270–15275.

61. Huang YW, Opriessnig T, Halbur PG, & Meng XJ (2007) Initiation at the third in-frame AUG codon of open reading frame 3 of the hepatitis E virus is essential for viral infectivity in vivo. J Virol 81(6):3018–3026.

62. Pudupakam RS, et al. (2009) Deletions of the hypervariable region (HVR) in open reading frame 1 of hepatitis E virus do not abolish virus infectivity: evidence for attenuation of HVR deletion mutants in vivo. Journal of virology 83(1):384–395.

63. Kenney SP (2019) The Current Host Range of Hepatitis E Viruses. Viruses 11(5).

64. Cossaboom CM, et al. (2012) Cross-species infection of pigs with a novel rabbit, but not rat, strain of hepatitis E virus isolated in the United States. The Journal of general virology 93Pt 8):1687–1695.

65. Johne R, et al. (2010) Novel hepatitis E virus genotype in Norway rats, Germany. Emerging infectious diseases 16(9):1452–1455.

66. Johne R, et al. (2010) Detection of a novel hepatitis E-like virus in faeces of wild rats using a nested broad-spectrum RT-PCR. The Journal of general virology 91(Pt 3):750–758.

67. Li TC, et al. (2011) Characterization of self-assembled virus-like particles of rat hepatitis E virus generated by recombinant baculoviruses. J Gen Virol 92(Pt 12):2830–2837.

68. Johne R, et al. (2012) Rat hepatitis E virus: geographical clustering within Germany and serological detection in wild Norway rats (Rattus norvegicus). Infection, genetics and evolution 12(5):947–956.

69. Nan Y & Zhang YJ (2016) Molecular Biology and Infection of Hepatitis E Virus. Front Microbiol 7:1419.

70. Tian D, et al. (2022) Hepatitis E virus infects brain microvascular endothelial cells, crosses the blood-brain barrier, and invades the central nervous system. Proc Natl Acad Sci U S A 119(24):e2201862119.

71. Halbur PG, et al. (2001) Comparative pathogenesis of infection of pigs with hepatitis E viruses recovered from a pig and a human. Journal of clinical microbiology 39(3):918–923.

72. Gojmerac T, Kartal B, Žurić M, Ćurić S, & Mitak M (1995) Serum biochemical and histopathological changes related to the hepatic function in pigs following atrazine treatment. Journal of Applied Toxicology 15(3):233–236.

73. Rico AG, Braun JP, Benard P, & Thouvenot JP (1977) Tissue and blood gamma-glutamyl transferase distribution in the pig. Research in Veterinary Science 23(3):395–396.

74. Emerson SU, et al. (2010) Release of genotype 1 hepatitis E virus from cultured hepatoma and polarized intestinal cells depends on open reading frame 3 protein and requires an intact PXXP motif. Journal of virology 84(18):9059–9069.

75. Yadav KK, Boley PA, Fritts Z, & Kenney SP (2021) Ectopic Expression of Genotype 1 Hepatitis E Virus ORF4 Increases Genotype 3 HEV Viral Replication in Cell Culture. Viruses 13(1).

76. Meyer RC, Bohl EH, & Kohler EM (1964) PROCUREMENT AND MAINTENANCE OF GERM-FREE SEINE FOR MICROBIOLOGICAL INVESTIGATIONS. Applied microbiology 12(4):295–300.

77. Xu J, et al. (2022) The Cold-Adapted, Temperature-Sensitive SARS-CoV-2 Strain TS11 Is Attenuated in Syrian Hamsters and a Candidate Attenuated Vaccine. Viruses 15(1).

78. Gupta P, et al. (2012) Immunohistochemistry for the diagnosis of hepatitis E virus infection. J Viral Hepat 19(2):e177–183.

